# A Comprehensive Epidermal Map from a Poplar Single-Cell Shoot Atlas Reveals New Trichome-Specific Genes

**DOI:** 10.64898/2026.07.02.736106

**Authors:** Anita Giabardo, Joshua C. Wood, Saurabh P. Pandey, Julia Brose, Sierra S. Cloud, John P. Hamilton, Alex D. Heise, Rakesh Loya, Ziliang Luo, Kathrine Mailloux, Brieanne Vaillancourt, Daniela L. Weber Wyneken, Robert J. Schmitz, Breeanna R. Urbanowicz, Chung-Jui Tsai, C. Robin Buell

## Abstract

Poplar (*Populus* spp.) is a model system for tree biology. Specifically, *P. tremula* x *P. alba* INRA 717-1B4 (hereafter “poplar 717”) has become an important platform for functional genomics and synthetic biology due to its rapid growth and ease of transgenesis. Here, we present a single-cell RNA-seq atlas of the poplar 717 shoot, including apical meristem, primary and secondary stems, and three stages of leaf development. Analysis of ca. 159,000 cells resolved 40 transcriptionally distinct clusters representing 7 major cell types, providing a high-resolution view of shoot development and tissue organization. We focused on the epidermis which constituted >15% of cells in the shoot atlas for in-depth characterization of epidermal heterogeneity. By integrating known marker genes with transcriptomic signatures consistent with established poplar leaf phytochemistry, we annotated epidermal cell subclusters corresponding to developmental stages, spatial location, and specialized cell types, including a distinct population of non-glandular trichomes. Coupling the single-cell RNA-seq atlas with bulk transcriptome data from glabrous mutants enabled the identification of novel trichome markers. Experimental validation of a representative trichome-specific promoter established a tool with potential to support cell type-targeted-metabolic engineering. We provide the poplar 717 atlas to the community through the BioPoplar Atlas Viewer (http://bio-poplar-atlas.com), providing a platform to explore the poplar transcriptome at single-cell resolution and a foundation for data-driven cell type-aware genetic engineering strategies in poplar.

## Introduction

Poplar (*Populus* spp.) is a well-established model system for tree physiology and bioenergy, bioproduct, and biomaterial production due to its fast growth, high biomass accumulation, low input requirements, and widespread geographical distribution (Sannigrahi, Ragauskas, and Tuskan 2010; Porth and El-Kassaby 2015; Joshi, Difazio, and Kole 2011). The interspecies hybrid *P. tremula* x *P. alba* INRA 717-1B4 (hereafter “poplar 717”) is particularly amenable to *Agrobacterium*-mediated transformation, regeneration (Leple et al. 1992) and genome editing (X. Zhou et al. 2015). The recent release of a haplotype-resolved genome (R. Zhou et al. 2023) has further advanced its use in functional genomics, gene editing, and synthetic biology research (Feyissa et al. 2025). To avoid negative pleiotropic effects due to over- or mis-expression, a detailed understanding of the spatiotemporal regulation of gene expression at the single-cell resolution is essential to enable data-driven cell type-aware genetic engineering strategies (Bibik et al. 2024).

Recent advances in single-cell genomic technologies now make it possible to interrogate the plant transcriptome at true cell-type resolution (Islam et al. 2024). Both protoplast and nuclei-based approaches have been successfully applied to an increasingly broad range of species, including the model species *Arabidopsis thaliana* (Lee et al. 2025), important food crops such as rice (X. Wang et al. 2025; Yan et al. 2025), soybean (*Glycine max*, Zhang et al. 2025), and maize (*Zea mays,* Marand et al. 2021), and medicinal species such as *Catharanthus roseus* (C. Li et al. 2023). In *Populus*, single-cell or single-nuclei RNA-seq datasets have been generated for stems (Tung et al. 2023; R. Li et al. 2023; H. Li et al. 2021; Jianbo Xie et al. 2022; Du et al. 2023; Y. Chen et al. 2021; Gómez-Soto et al. 2025; Schmidt et al. 2025), leaves (Jing et al. 2024), and shoot apical meristem (SAM) (Conde et al. 2022). These efforts span multiple *Populus* species, a range of developmental stages, plant growth conditions, and single-cell technology platforms. Currently, poplar 717 is only represented by single-nuclei-based SAM (Conde et al. 2022) and stem (Gómez-Soto et al. 2025) datasets. Leaf tissues are surprisingly underrepresented, despite their central role as energetic hubs and their capacity for chemically and functionally diverse specialized metabolism, such as flavonoids, terpenoids, and phenolics across distinct leaf cell types (Weng et al. 2021; Sun et al. 2023). Beyond their importance in defense, signaling, and environmental adaptation, many of these complex molecules represent additional opportunities for metabolic engineering, complementing efforts focused on lignocellulosic biomass and secondary cell wall (SCW) biology.

To further develop poplar as a multipurpose feedstock crop for biomaterials and bioproducts (Buell et al. 2023), we generated a comprehensive, protoplast-based single-cell transcriptome atlas of poplar 717 across shoot development, including the SAM, three leaf stages, and two stem stages. Given the importance of poplar as a source of lignocellulosic biomass, we analyzed the expression profiles of genes involved in SCW biosynthesis across the atlas, identifying two xylem clusters and one epidermal cluster enriched with SCW-related genes. In the poplar shoot, the epidermis is present in all organs, serving diverse functions including protection from UV light, mechanical stress, regulation of water loss and gas exchange, and temperature control through specialized features such as the cuticle, trichomes, and stomata. We annotated 13 epidermal subclusters representing all major epidermal cell types, including abaxial and adaxial pavement, stomata, hydathode, and trichome cells. We functionally validated trichome subclusters providing a foundation for targeted metabolic engineering of poplar. The poplar 717 shoot single-cell atlas is publicly available through the BioPoplar Atlas Viewer (http://bio-poplar-atlas.com), providing an accessible and high-resolution resource for the research community.

## Results

### A comprehensive single-cell shoot atlas of poplar 717

Using the microfluidics-free Particle-templated Instant Partitioning sequencing (PIP-seq) technology (Clark et al. 2023), we generated a single-cell RNA-seq (scRNA-seq) atlas using protoplasts from six tissues: SAM, three leaf stages (L1, L3, and L5), and primary (PS) and secondary (SS) stems (Fig. 1a). Following optimization of protoplast isolation, each organ and developmental stage was processed separately. Reads were mapped to the haplotype-resolved 717 genome (R. Zhou et al. 2023) and replicates integrated at the organ level. We then applied Uniform Manifold Approximation and Projection (UMAP) to determine the representation across the organs. Altogether, this atlas includes 159,353 cells and captures 94% of the total genes present in the two 717 haplotypes (Fig. 1b) (R. Zhou et al. 2023). The number of cells, number of genes detected, and number of Unique Molecular Identifiers (UMIs) varied depending on organ and developmental stage. The numbers ranged between 12,119 and 47,006 cells per dataset (Fig. 1c, upper). The total number of genes detected in each dataset was comparable across organs (Fig. 1c, lower), suggesting adequate detection of the transcriptome. A set of 50,741 genes (84%) was detected in all organs and developmental stages. L1, L3 and L5 shared expression of 52,500 genes, yet distinct subsets of genes were detected in only one or two developmental stages (Fig. 1d, left). Among the two stem developmental stages, 53,706 genes were detected in both PS and SS, with 2,040 specific to PS and 1,406 specific to SS (Fig. 1d, right).

**Fig. 1:**
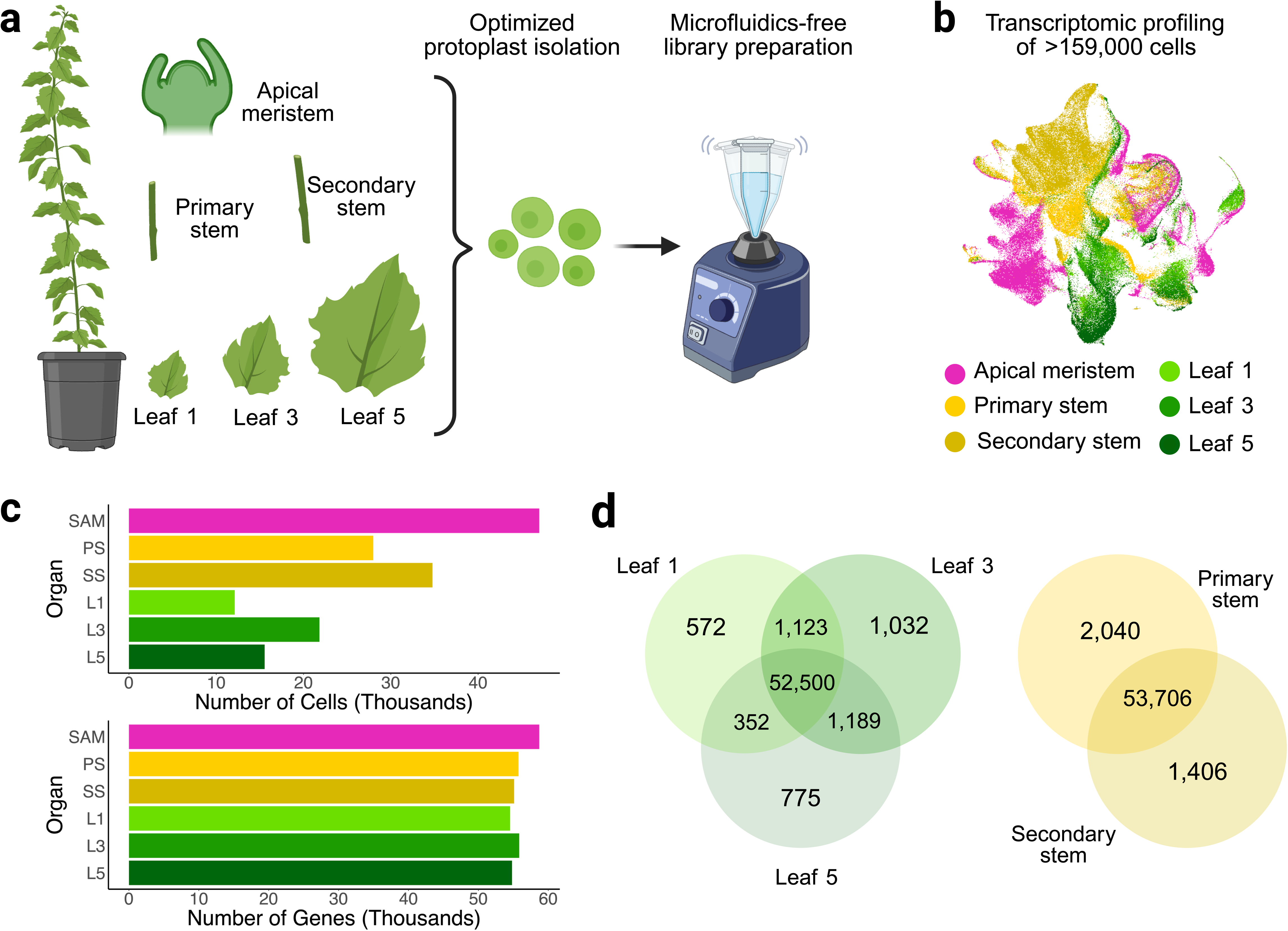
Generation of the scRNA-seq poplar shoot atlas. **a**) Tissues from growth-chamber grown hybrid poplar 717 were sampled. For each tissue, protoplast isolation was carried out using an optimized protocol, followed by PIP-seq library preparation via vortexing. **b**) Uniform Manifold Approximation and Projection (UMAP) plot of >159,000 cells composing the scRNA-seq poplar shoot atlas, colored by organ. **c**) Upper: number of cells profiled in each organ. Lower: number of genes detected in each organ. SAM: shoot apical meristem; PS: primary stem; SS: secondary stem; L1: Leaf 1; L3: Leaf 3; L5: Leaf 5. **d**) Venn diagrams showing the number of genes shared across different developmental stages within the leaf and stem series. Created in BioRender.

To assess changes in cell-type composition and abundance across organs and developmental stages, we integrated replicate and developmental datasets at the organ level (Fig. 2). We identified 25, 23, and 20 UMAP clusters for SAM (Fig. 2a), stem (Fig. 2b), and leaf (Fig. 2c), respectively. Chi-Squared analyses revealed no significant differences in cluster composition between biological replicates. Conversely, multiple clusters showed over- or under-representation across developmental stage. For example, Cluster PS_11 (Fig. 2b, left, arrowheads) was significantly enriched relative to SS_11 (Fig. 2b, right, arrowheads; 2,311 cells vs. 11 cells, p < 0.01), whereas the opposite pattern was observed for Cluster PS_6 and SS_6 (146 cells in PS vs. 2,977 cells in SS, p < 0.01; Fig. 2b, arrowheads). In leaves, a significant progressive decrease in the number of cells (p < 0.01) was observed for Cluster 4 in development from L1 to L5 (1,963 cells in L1 vs. 143 cells in L5) and Cluster 19 in development from L1 to L5 (95 cells in L1 vs. 31 cells in L5), while Cluster 0 displayed the opposite trend (Fig. 2c, arrowheads). These results suggest that integrated analyses can reveal developmentally regulated changes in cell-type composition.

**Fig. 2:**
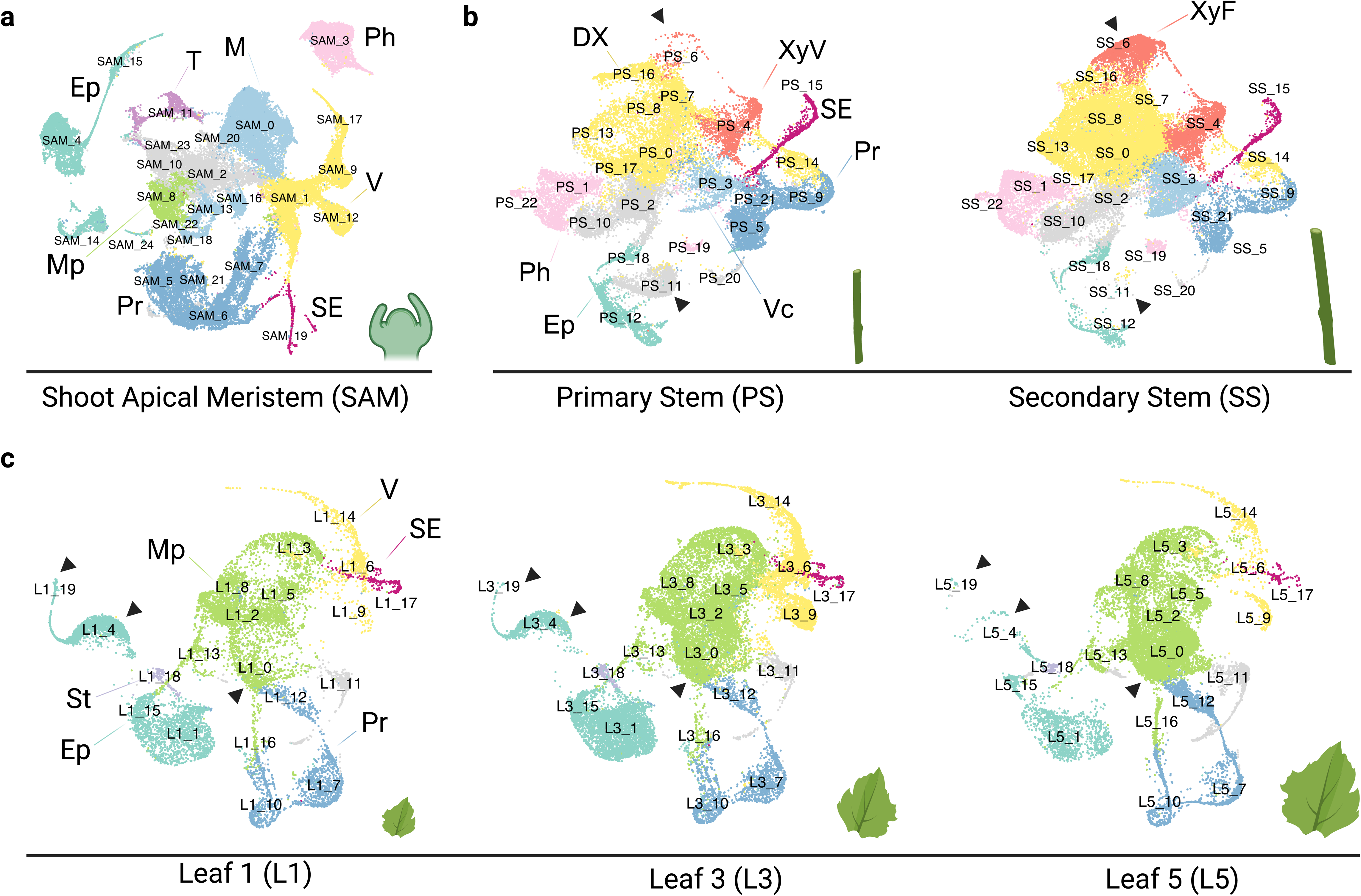
UMAP plots of organ-level data. **a**) UMAP of the shoot apical meristem dataset. **b**) UMAP of the integrated primary and secondary stem datasets, split by developmental stage. **c**) UMAP of the integrated leaf datasets (including L1, L3, and L5), split by developmental stage. Main cell types identified are labeled as follows. Ep: epidermis; DX: developing xylem; Mp: mesophyll; Ph: phloem; Pr: proliferating; St: stomata; SE: sieve elements; T: trichomes; V: vasculature; Vc: vascular cambium; XyV: xylem vessel; XyF: xylem fiber. Arrowheads indicate clusters referenced in the text. Created in BioRender.

We then merged all datasets to generate a more comprehensive shoot atlas. UMAP analysis resulted in 40 distinct clusters (Fig. 3a). Cluster annotation for both cell cycle (Fig. 3c, upper) and cell type (Fig. 3c, lower) was performed via known marker genes curated from the literature, including previous poplar (Conde et al. 2022; Daniela Gómez-Soto et al. 2025) and *A. thaliana* scRNA-seq studies (Tenorio Berrío et al. 2022; J.-Y. Kim et al. 2021; Xia et al. 2022; Lee et al. 2025; Schmidt et al. 2025). UMAP clusters were grouped into major, color-coded cell states and tissue identities, including meristematic and proliferating cells, as well as differentiated epidermal, ground, mesophyll, xylem, and phloem lineages (Fig. 3a). The majority of clusters contained cells from all sampled organs (Fig. 3b), suggesting that clustering was driven primarily by cell type and state rather than organ of origin. Notable exceptions included multiple meristematic clusters composed predominantly of SAM cells (Clusters 1, 9, 10, 17, 29, and 35), xylem and phloem clusters derived mainly from SS cells (Clusters 5, 21, 23, 13, and 16), and mesophyll and ground tissue clusters dominated by cells from leaves (Clusters 30, 0, 32) (Fig. 3b, lower). These patterns likely reflect either specialized cell states or specialized cell types. Nevertheless, all major groups in the shoot atlas captured clusters derived from multiple organs, providing an opportunity to investigate developmental regulation of closely related cell types across organs. Using the 3,000 most variable genes across the atlas, correlation of gene expression recapitulated cell-type annotation.

**Fig. 3:**
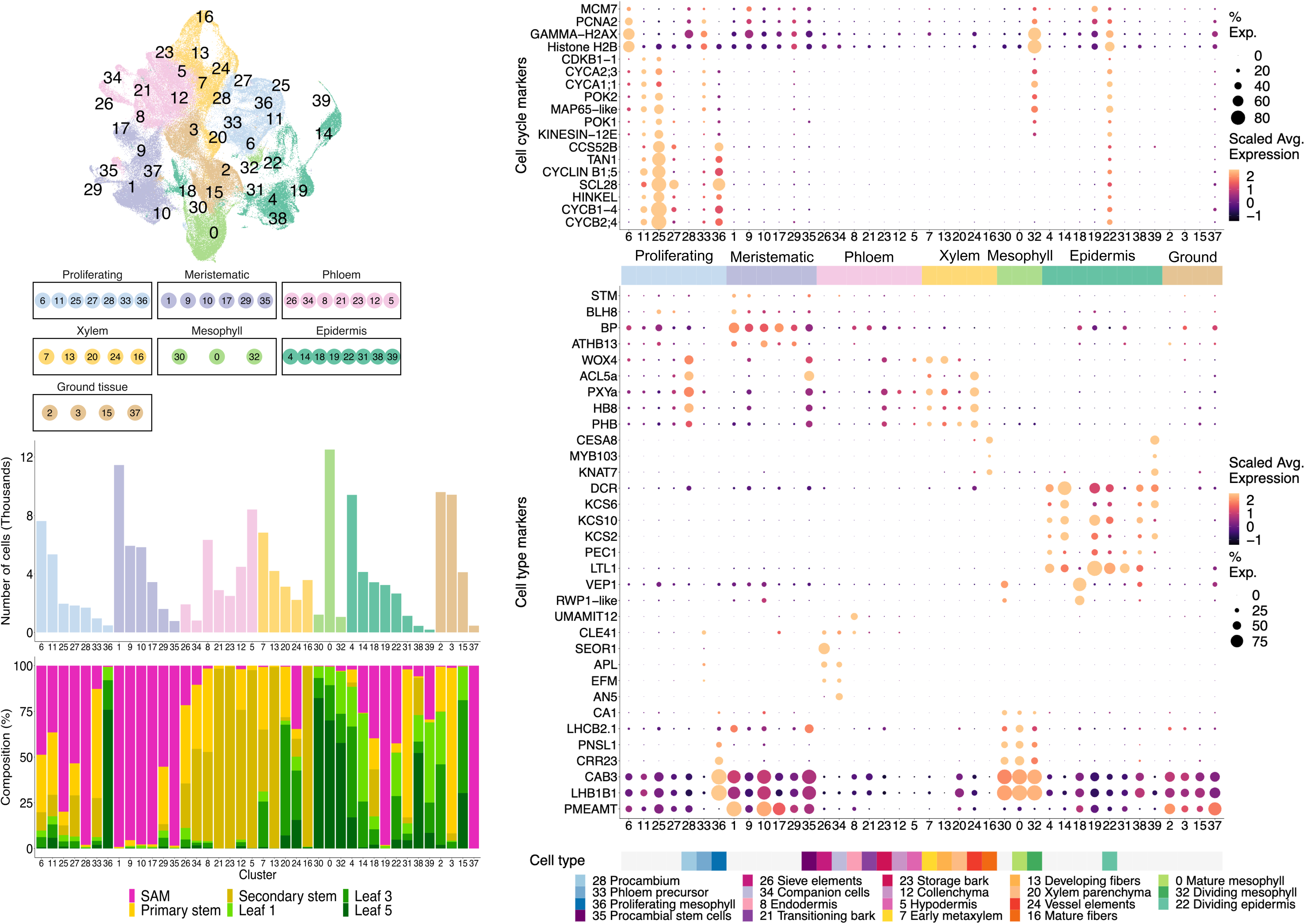
Overview and annotation of the scRNA-seq poplar shoot atlas. A) UMAP plot (upper) including all cells profiled, after integration and filtering, colored by cell type as indicated by the legend (lower); B) Upper: number of cells in each cluster, colored by cell type as indicated in A; Lower: share of cells belonging to the different tissues included in the atlas. C) Upper: dot plot showing cell cycle markers. Lower: dot plot showing cell type markers. Stars denote genes previously validated via in-situ hybridization or promoter:reporter lines. Created in BioRender.

Based on established cell cycle markers (Fig. 3c, upper), Clusters 6, 11, 25, 27, 28, 33, and 36 were annotated as proliferating cells. While many of these clusters were composed of a high proportion of cells from the SAM (Clusters 6, 25, 27, and 28), others were enriched for stem (Cluster 33) and leaf (Cluster 36) cells. Based on additional cell type markers, we further annotated Cluster 28 as procambial cells, Cluster 33 as phloem precursors, and Cluster 36 as proliferating mesophyll cells (Fig. 3c, bottom). The detection of cell proliferation markers in SAM and procambium is consistent with prior scRNA-seq in poplar SAM and stem (Conde et al. 2022; Schmidt et al. 2025). Residual proliferation signals were observed in epidermal Cluster 22 and mesophyll Cluster 32, both of which were adjacent to proliferation cell clusters on the UMAP (Fig. 3a). Furthermore, the *cellulose synthase-like D5* (*CSLD5*) genes (Suzuki et al. 2006), recently shown to be involved in plate formation during cell proliferation (Lan et al. 2025), were specifically expressed in Clusters 11, 25, 33, 36, and 22 (Fig. 4).

**Fig. 4:**
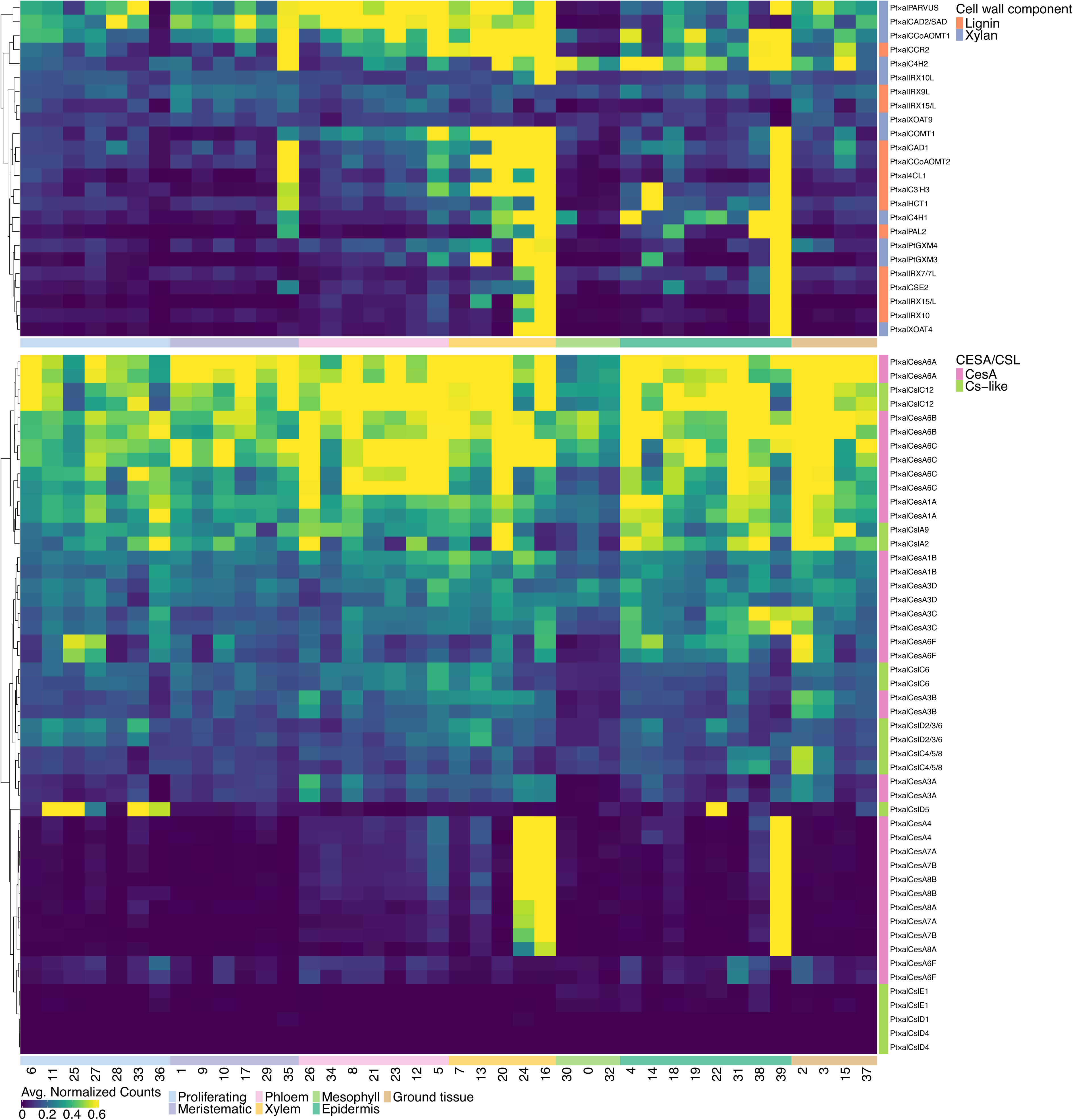
Expression of cell wall biosynthesis genes across the shoot atlas with lignin and xylan biosynthetic genes in the top panel and cellulose-related genes in the bottom panel. In the top panel, only genes from the *P. tremula* haplotype with average expression > 0.1 are shown. Expression (average normalized counts) of all genes is capped at the 90^th^ percentile.

### Expression of secondary cell wall biosynthesis genes is limited to specific xylem and epidermis clusters

Genes involved in the biosynthesis of the major SCW polymers cellulose, xylan and lignin were well-represented in xylem Clusters 16 and 24 (Fig. 4). These include SCW-associated cellulose synthase genes *CESA8, CESA4* and *CESA7* (D. Song, Shen, and Li 2010), core xylan biosynthesis and side-chain modification genes (Zhong, Cui, and Ye 2019), lignin pathway genes, and multiple upstream master regulators of SCW transcriptional networks (Zhong, Cui, and Ye 2019). Among the highest-ranking transcripts in Cluster 16 were the lignin biosynthetic genes *Caffeic Acid O-Methyltransferase 1* (*COMT1*, 3.40 average normalized counts) and *Caffeoyl-CoA O-Methyltransferase 1* (*CCoAOMT1*, 2.58 average normalized counts) as well as the xylan-associated *UDP-xylose synthases 6* (*UXS6) and UXS5.* Cluster 16 consisted almost entirely of cells from the SS (Fig. 3b), consistent with a cell population actively engaged in SCW biogenesis. Notably, the DUF579 family gene *DUF579-9* (Potri.005G141300), which functions in glucuronoxylan biosynthesis, exhibits fiber-specific promoter activity in poplar and has been used for fiber-targeted cell wall engineering (Gui et al. 2020; J. Li et al. 2025). These features led us to annotate Cluster 16 as xylem fiber cells.

In contrast, Cluster 24 contained cells from all six organs (Fig. 3) with lower SCW gene expression relative to Cluster 16 (Fig. 4). Genes more abundant in Cluster 24 included the vessel marker *MAN6* (Potri.016G138600) and its genome duplicate *MAN4* (Potri.006G109900), encoding endo-mannanase (Zhao et al. 2013), as well as multiple *XCP* genes encoding xylem cysteine proteases. Although XCPs are frequently considered programmed cell death (PCD) markers due to their function in autolysis during xylogenesis, they are synthesized as inactive precursors and stored in the vacuole prior to initiation of PCD (Funk et al. 2002; Avci et al. 2008). Consistent with this developmental role, the poplar *XCP1* (Potri.004G207600) promoter exhibits vessel-specific expression and has been used for vessel-targeted cell wall engineering (Gui et al. 2020).Thus, we annotated Cluster 24 as vessel cells containing a developmental continuum in contrast to the more uniformly SCW-active fiber population in Cluster 16. This annotation is further reinforced by the differential enrichment of vessel-specific master regulator wood-associated NAC domain transcription factor genes *WND5A* and *WND5B* in Cluster 24 and the fiber-specific *WND1A* and *WND1B* in Cluster 16 (Zhong et al. 2008; Zhong, Lee, and Ye 2010).

Cluster 39, annotated as epidermis, was also enriched for SCW-related genes (Fig. 4) and composed primarily of cells from the SAM (29%) and L1 (44%) (Fig. 3b). This cluster contained *MYB183a*, one of the eight poplar 717 genes encoding trichome initiation regulators (Bewg et al. 2022). Cluster 39 showed high correlation with Cluster 14 (0.62), in which all eight trichome-regulating *MYB* genes were highly expressed. Based on its SCW gene expression, cluster composition, and its relatively small size (187 cells, the smallest cluster in our atlas), we hypothesize that Cluster 39 represents developing trichome cells, whereas the larger neighboring Cluster 14 likely represents trichome initials.

### Annotation of shoot epidermis reveals epidermal cell functional diversification

All of the organs included in the atlas have epidermis thereby providing an opportunity to understand epidermis cell diversity across different organs and development. Thus, we re-clustered the 24,011 epidermal cells from the full ∼159,000 atlas (Clusters 4, 14, 18, 19, 22, 31, 38, 39; Fig 3a) to generate an epidermis-only UMAP with 14 subclusters (Fig. 5a; labelled Subcluster epi_#).

**Fig. 5:**
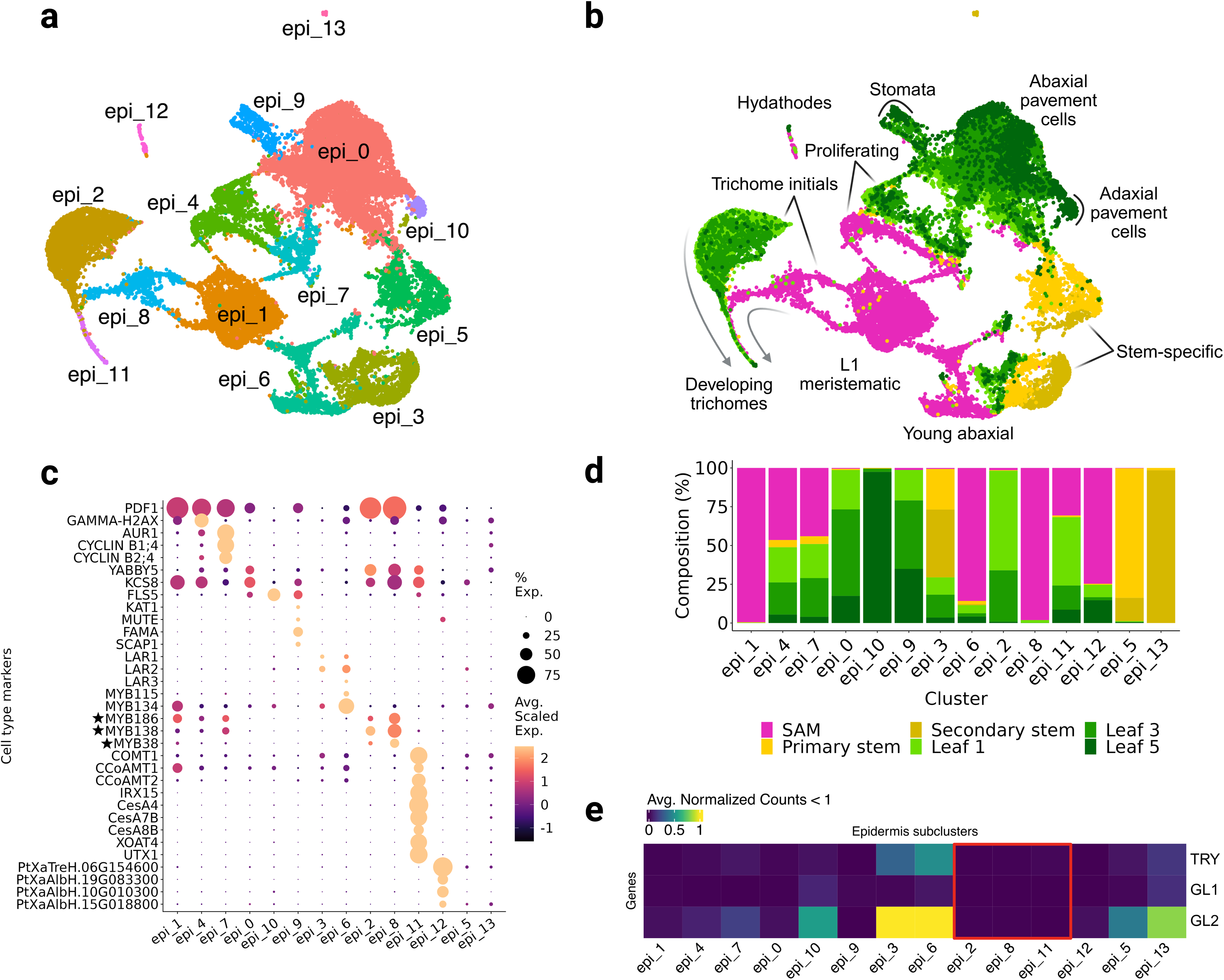
Annotation of the epidermis map. **a**) UMAP analysis reveals 14 distinct epidermis subclusters. **b**) Epidermis UMAP colored by tissue of origin. **c**) Expression of marker genes used to annotate the epidermis subclusters. Stars indicate experimentally validated trichome initiation regulators in poplar (Bewg et al. 2022). **d**) Composition of epidermis subclusters. **e**) Arabidopsis genes previously used for trichome annotation are not expressed in the putative trichome subclusters (boxed in red). Created in BioRender.

Subcluster epi_1 was constituted predominantly of SAM cells with high expression of *PROTODERMAL FACTOR 1* (*PDF1*), specific to the L1 layer and protoderm (Abe, Takahashi, and Komeda 1999) likely representing L1 cells of the meristem. Top differentially expressed genes in Subcluster epi_4 included known cell cycle progression markers, including S-phase histone proteins such as histone *H2A* genes and *GAMMA-H2AX* (Fan et al. 2022) (Fig. 5c). Subcluster epi_7 was enriched in cell cycle progression markers including *ATAURORA 1* (*AUR1*) (Demidov et al. 2005), *CYCLIN B1;4*, and *CYCLIN B2;4* (Vandepoele et al. 2002). Both Subclusters were annotated as proliferating cells.

We annotated epi_0 as abaxial pavement cells based on strong expression of abaxial marker *YABBY5* (Sarojam et al. 2010) and multiple *KCS* (*3-ketoacyl-CoA synthase*) genes involved in very long chain fatty acid biosynthesis that form the backbone of cuticular wax (Gonzales-Vigil et al. 2017). This Subcluster also showed expression of flavonol branch pathway genes such as *FLS5* (*flavonol synthase*), but not *DFR* (*dihydroflavonol reductase*) and other downstream genes required for anthocyanin and proanthocyanidin biosynthesis (Tsai et al. 2006). The adjacent, small epi_10 Subcluster, enriched for L5 cells, showed higher expression of flavonol branch pathway genes than epi_0. Flavonols such as kaempferol, quercetin and their sugar conjugates are routinely detected in poplar leaves and serve a UV-protective function (H.-Y. Chen et al. 2014); thus, epi_10 likely represents more mature adaxial pavement cells. The marker genes *FAMA, SCAP1* (*STOMATAL CARPENTER 1*)*, MUTE* (Lopez-Anido et al. 2021) and *KAT1* (*3-KETO-ACYL-COA THIOLASE 1*) (Ichida et al. 1997) were highly expressed in epi_9, suggesting this Subcluster represents guard cells and stomatal lineage cells.

Proanthocyanidins (condensed tannins) accumulate in abaxial epidermis of poplar leaves (Kao, Harding, and Tsai 2002). Epi_3 and epi_6 expressed both early and late-stage flavonoid pathway genes such as *anthocyanidin reductases* (*ANRs*) and *leucoanthocyanidin reductases* (*LARs*), which are specifically involved in condensed tannins biosynthesis (Tsai et al. 2006). Compared to epi_3, epi_6 appeared younger based on a greater enrichment of cells from SAM and higher expression of ribosomal and histone genes as well as higher expression of condensed tannins master regulators *MYB115* and *MYB134* (James et al. 2017). Therefore, we annotated epi_6 as young abaxial epidermis and epi_3 as expanding abaxial leaf epidermis and stem epidermis.

Epi_5, mainly constituted of PS and SS cells, was annotated as stem-specific epidermis. The small epi_12 and epi_13 Subclusters did not express any known cell-type-specific poplar markers. Based on high expression of genes orthologous to experimentally verified *Arabidopsis* hydathode-specific chitinases, AT5G24090, AT3G54420, and AT4G19810, and an extensin-like gene AT3G16660 (Yagi et al. 2021), epi_12 likely represents hydathode cells. Epi_13 may represent contamination with photosynthesizing mesophyll or older stem epidermis.

Epi_2 and epi_8 Subclusters shared 14 out of their top 20 differentially expressed genes (log2 FC > 2, adjusted p-value < 0.05). Known regulators of trichome initiation *MYB186*, *MYB138*, and *MYB38,* were expressed in both epi_2 and epi_8, but were lowly expressed in the much smaller epi_11 Subcluster. By contrast, genes involved in SCW deposition and remodeling were highly expressed in Subcluster epi_11, including multiple lignin genes (*COMT1*, *CCoAMT1*, and *CCOAMT2*), as well as SCW-associated *CesA4*, *CesA7*, and *CesA8* (Fig. 5c). Gene ontology (GO) analysis of the top differentially expressed genes (log2 FC > 2, adjusted p-value < 0.05) in each subcluster revealed enrichment (adjusted p-value < 0.05) in cell wall-related terms (“xylan biosynthetic process”, “cellulose biosynthetic process”) in Subcluster epi_11. Consistent with these results, additional genes with peak expression in epi_11 included orthologs of *Arabidopsis* xylan biosynthetic genes, such as XOAT4 (also known as *TRICHOME BIREFRINGENCE-LIKE 3*), an *O*-acetyl transferases, *IRX15-LIKE* (Smith et al. 2017), and *UDP-XYLOSE TRANSPORTER 1* (*UTX1*), which encode a Golgi-localized transporter required for UDP-xylose import during xylan matrix polysaccharide biosynthesis (Smith et al. 2017; Ebert et al. 2015) (Fig. 4c).

Based on the expression of known trichome regulators and SCW genes, and the composition of Subclusters epi_2 (3,062 cells, 64% L1), epi_8 (895 cells, 98% SAM), and epi_11 (186 cells, 30% SAM, 44% L1, 15% L3), we infer that these three Subclusters represent a trichome development gradient. Cells in epi_2 and epi_8 represent trichome initials in leaves and the SAM, respectively, whereas cells in epi_11 represent developing trichomes with active SCW gene expression.

### Identification of Trichome-Specific Markers Using Glabrous Poplar Mutants

Previously published poplar single-nuclei RNA-seq data inferred trichome identity primarily using orthologs of the *Arabidopsis* genes *TRY* (Potri.015G022000), *GL1* (Potri.012G080400), and *GL2* (Potri.003G052400) (Conde et al. 2022). In *Arabidopsis*, however, *TRY*, *GL1* and *GL2* are components of a broader epidermal patterning network whose outputs include, but are not limited to, trichome formation (Serna 2005; S.-K. Song et al. 2024; S. Wang et al. 2010). In this study, the *GL1* ortholog was poorly expressed, while *TRY* and *GL2* showed highest expression in shoot Clusters 18 and 38 and in epidermal Subclusters epi_3 and epi_6 (Fig. 5e), which we annotated as abaxial epidermal cells (Fig. 5b-c). These genes were absent from the trichome-associated Subclusters epi_2, epi_8, and epi_11, which were instead defined by expression of experimentally validated trichome initiation regulators in poplar (Plett et al. 2010; Bewg et al. 2022).

To resolve this discrepancy and to identify additional trichome markers independent of epidermal fate, we leveraged the glabrous 717 mutants (Fig. 6a) generated by CRISPR multiplex editing of *MYB186*, *MYB138* and *MYB83* (Bewg et al. 2022). Bulk RNA-seq of L1 tissues from three WT plants and three independent glabrous mutant lines identified 592 differentially expressed genes (adjusted p-value < 0.01, |log_2_FC| > 2, Fig. 6b). The vast majority (582) were downregulated in the trichomeless mutants, including the targeted *MYB* regulators themselves. These genes showed enrichment for SCW biogenesis, lipid transport, and triterpenoid biosynthesis, consistent with established roles of non-glandular trichomes in plant surface defense (Bewg et al. 2022; Karabourniotis et al. 2020). We reasoned that genes downregulated in the glabrous mutants may exhibit trichome-specific expression. Of the top 29 most strongly downregulated genes (log_2_FC < -10), three were not detected and twelve were expressed at very low levels (< 0.2 average normalized counts) across epidermis subclusters. The remaining 14 genes were exclusively detected in three annotated trichome subclusters of epidermis: six in all trichome subclusters, four–including *MYB38*–in trichome initials, and five in developing trichomes (Fig. 6c-d). Trichome-specific expression was also observed in the shoot atlas, with the exception of *LAC3* (*laccase*) which was also expressed in the xylem fiber Cluster 16, consistent with its role in lignin polymerization and SCW formation (Ranocha et al. 2002).

**Fig. 6:**
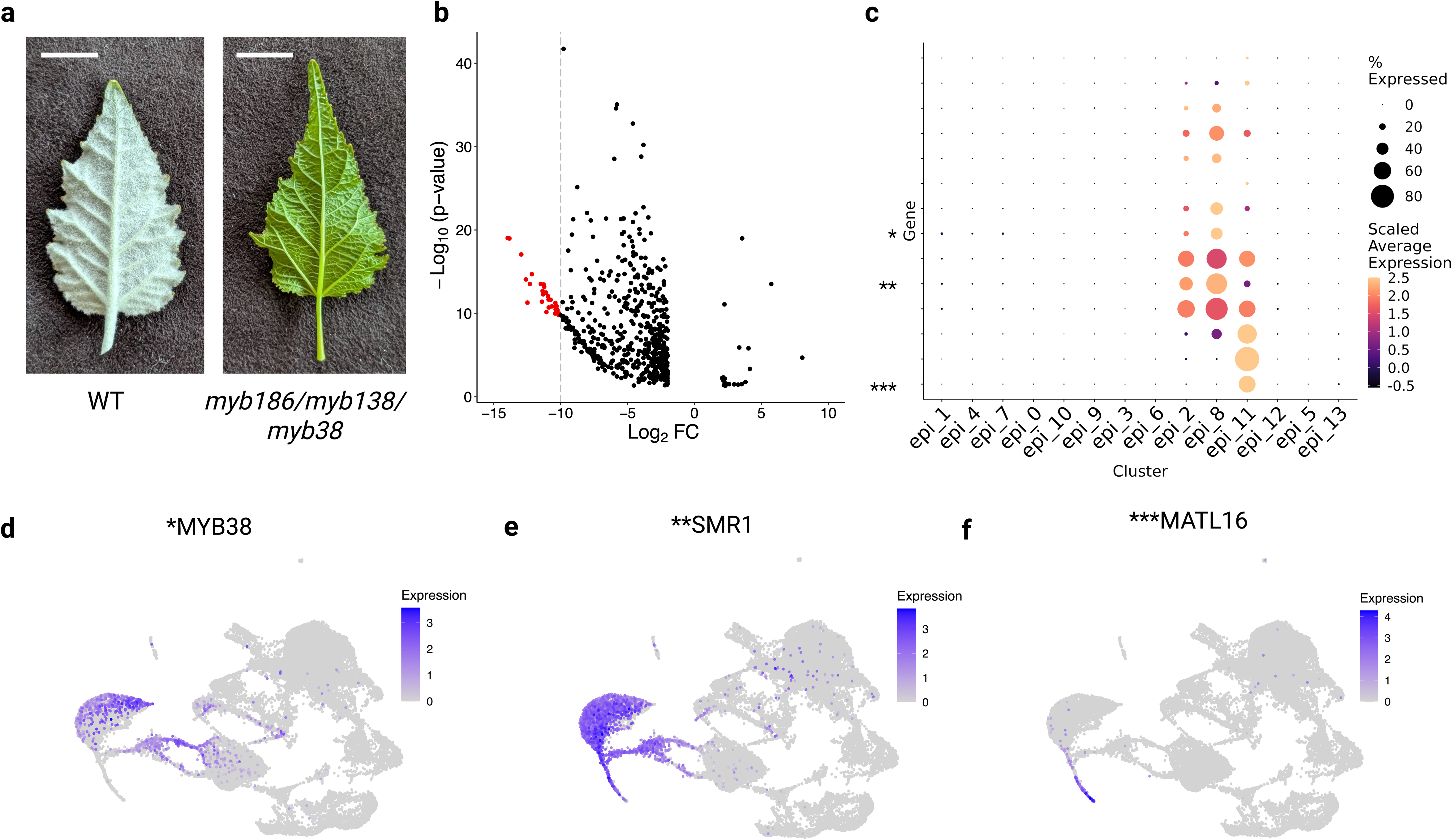
Integration with bulk RNA-seq differential gene expression analysis validates putative trichome cluster annotation. **a**) Pictures of abaxial leaf 2 in WT (left) and representative glabrous mutant (right) poplar 717. Scale bars: 1 cm. **b**) Volcano plot showing differentially expressed genes. Genes in red (log2FC < -10, p-value < 0.05) were selected for further analysis. **c**) Expression profile of 14 differentially expressed genes selected in b. Only genes with average expression >0.2 in at least one epidermis subcluster are shown. Asterisks refer to the genes shown in panel d-f. **d**-**f**) Expression of genes selected from c. Created in BioRender.

Genes detected in all three trichome subclusters included orthologs implicated in lipid metabolism, such as fatty acid desaturase and diacylglycerol acyltransferase, as well as both alleles of poplar *SMR1* (Churchman et al. 2006), which encode a SIAMESE-like (SMR) cyclin-dependent protein kinase inhibitor (Fig. 6e). SMRs regulate the trichome endocycle by promoting endoreduplication, and this function is broadly conserved in land plants (Kumar et al. 2015; Walker et al. 2000). Their widespread expression across all three trichome subclusters is therefore consistent with the active endoreduplication program that characterizes non-glandular trichome development (Walker *et al*., 2000). This pattern also supports our inference that these clusters represent a developmental continuum rather than discrete cell types (Fig 6e).

Genes expressed specifically in the developing trichome Subcluster epi_11 included *LAC3* which is involved in lignin polymerization (Ranocha *et al*., 2002) and a suberization-associated *MYB59* (Lashbrooke et al. 2016; Rains, Gardiyehewa de Silva, and Molina 2018). The MYB59 orthologs in *Arabidopsis*, AtMYB9 and AtMYB107, were recently shown to regulate lignification and suberization of the seed coat (Hyvärinen et al. 2025). These findings are consistent with the enrichment of lignification and lipid-related genes in the trichome subclusters. The most strongly expressed trichome markers were two alleles of *PtaMATL16* (PtXaAlbH.10G166200 and PtXaTreH.10G173100, 2.3-2.6 average normalized counts), a poplar-specific malonyltransferase-like gene of the BAHD acyltransferase family (Tuominen, Johnson, and Tsai 2011). Both *MATL16* alleles were detected exclusively in developing trichomes of epidermal Subcluster epi_11 and in shoot Cluster 39. To validate trichome specificity, we cloned a 1.8 kb *PtMATL16* (PtXaTreH.10G173100) promoter fragment to drive expression of the free-use green fluorescent protein (fuGFP) gene (Moratti et al. 2024) in poplar 717. Confocal imaging of young leaves from multiple independent transgenic events detected PtMATL16::GFP activity exclusively in trichomes (Fig. 7), in contrast to CaMV35S::fuGFP which were found in both trichome and epidermal cells. These data establish *PtaMATL16* as a highly specific marker gene of developing non-glandular trichomes in poplar.

**Fig. 7:**
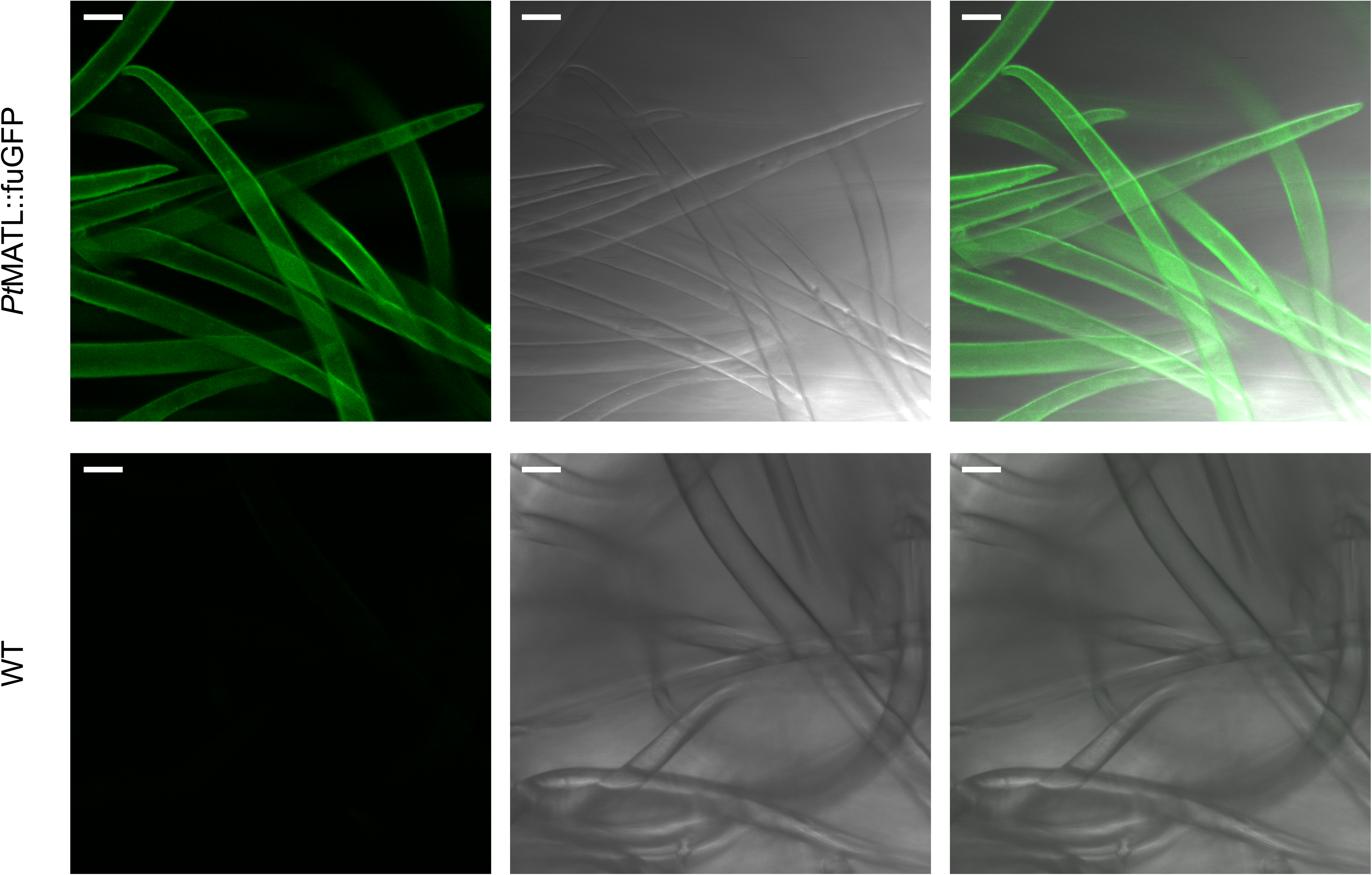
Trichome-specific GFP activity driven by the PtMATL16 promoter. Top row: Confocal and bright-field images from PtMATL16::fuGFP transgenic poplar showing strong GFP signals restricted to non-glandular trichomes. Bottom row: Corresponding images from wild-type leaves showing no detectable GFP fluorescence. Left panels show GFP channel, middle panels show bright field, and right panels present merged GFP/bright-field images. Scale bars, 20 µm.

### Construction of a Poplar Single-Cell Data-Mining Resource

To promote the use of the poplar 717 single-cell atlas and expand the *Populus* research community’s analytical resources, we developed the BioPoplar Atlas Viewer (http://bio-poplar-atlas.com/). This interactive platform allows users to seamlessly navigate through the processed and annotated scRNA-seq datasets generated in this study, providing valuable insights into gene expression across diverse poplar tissues, developmental stages and cell types. Prominent features of the BioPoplar Atlas Viewer include UMAP dimensionality-reduction plots that highlight distinct cell clusters along with heatmaps and dot plots that illustrate gene expression variation, facilitating in-depth comparative analyses. For instance, users can examine gene features within the UMAP plots to discover specific expression patterns, such as the enrichment of *PtMATL16* (PtXaTreH.10G173100) in trichomes (Cluster 39) in our leaf dataset. Furthermore, the BioPoplar Atlas Viewer enables users to visualize multiple genes of interest simultaneously, facilitating comparative analyses.

## Discussion

Hybrid poplar 717 is an important experimental system in tree biotechnology and functional genomics. In this study, we generated, annotated, and analyzed a comprehensive scRNA-seq atlas of the 717 shoot, including SAM, leaves, and stems. The atlas enabled the identification of SCW-depositing clusters and the annotation of epidermis subclusters with distinct functions across SAM, leaves, and stems. The comprehensive annotation of the epidermis revealed a trichome development gradient and novel trichome marker genes.

Poplar vascular tissues have been well characterized at the biochemical, molecular and genetic level, including single-cell/nuclei transcriptome analyses (R. Li et al. 2023; Schmidt et al. 2025). Here, we expand upon these studies and provide cell-type aware expression of genes involved in cellulose, lignin, and xylan biosynthesis in shoot development from SAM through three leaf stages and two stages of stem. Not only did we confirm elevated expression of lignin and xylan genes in developing xylem and xylem cells, but we also revealed diverse expression profiles of cellulose synthase and cellulose synthase-like genes across shoot development. We highlighted the specificity of *CESA4*, *CESA7,* and *CESA8* for secondary cell walls of xylem and trichomes, and the specificity of *CSLD5* for actively proliferating cells. Other *CesA* and *CSL* genes did not show specific expression patterns but instead were uniformly expressed across the entire atlas. Additionally, our data supports SCW deposition in poplar trichomes, consistent with the presence of lignin in *A. thaliana* trichomes (Marks et al. 2008) and in the recently discovered “neck strip” of glandular trichomes in cucumber (*Cucumis sativus*, Hao et al. 2024).

In poplar, the above-ground epidermis serves diverse functions including variable UV light incidence, pest and pathogen pressure, and water relations. As the epidermis is present in all organs, >15% of the cells captured in our shoot scRNA-seq atlas were epidermis which permitted a deep and robust annotation of epidermal cell types. Not surprisingly, clustering of epidermal cells across SAM, leaves and stems revealed some overlap in organ representation among the epidermis subclusters. With representation of leaf epidermis development series from SAM through to L5, we were able to annotate leaf abaxial and adaxial epidermis subclusters as well as highly specialized cells such as stomata, hydathodes, and trichomes. Data-mining these subclusters, including cell-type aware gene coexpression networks, can be a powerful strategy to investigate the impact of environmental perturbations on the epidermis.

The study of plant trichomes require optimized, species-specific isolation protocols (Marks et al. 2008; Huebbers et al. 2022) with bulk RNA-seq of trichome-enriched fractions as the primary strategy to study the trichome transcriptome (Sugimoto et al. 2022). Recently, scRNA-seq has proven useful for the study of glandular trichomes in *Nicotiana tabacum* (Chen et al. 2024). Poplar trichomes have been particularly elusive to study: they are morphologically different from *Arabidopsis* and *Nicotiana*, as they are non-glandular, unicellular, and non-branched (Plett et al. 2010). Although prior single-cell studies (Conde et al. 2022) used the *Arabidopsis* trichome development genes *TRY*, *GL1* and *GL2* (Szymanski et al. 1998) as markers, validated trichome-specific markers are lacking for poplar. *TRY* and *GL2* did not show specificity to either of the trichome clusters we identified. *TRY* and *GL2* did show cluster-specific expression in our SAM dataset, suggesting that the expression of *TRY* and *GL2* may be important for early stages of epidermal cell fate determination in the SAM but not directly involved in trichome development in poplar.

Through clustering of all epidermal cells from SAM, leaf, and stem coupled with differential gene expression analyses with WT and glabrous mutants, we identified a high-confidence set of 14 trichome marker genes (Fig. 6c) in poplar, supported by multiple independent lines of evidence. These markers span key stages of trichome development and include the experimentally validated trichome-initiation regulator *MYB38* (Bewg *et al*., 2022) enriched in trichome initials; the evolutionarily conserved endoreduplication regulator *SMR1* (Churchman *et al*., 2006) expressed across all trichome subclusters; and two markers specific to developing trichomes, including the lignin-associated *LAC3* (Ranocha et al. 2002) and the poplar-specific *MATL16* (Tuominen *et al*., 2011). Through promoter-report fusions, we validated *MATL16* as a trichome specific gene providing a new resource in metabolic engineering of trichome-specific bioproducts including biofuel precursors.

Non-glandular trichomes have long served as a model for single-cell differentiation and morphogenesis studies, most notably in *Arabidopsis* with its branched trichomes (Szymanski, Lloyd, and Marks 2000; Hülskamp 2004). The single-cell atlas presented here highlights both conserved and divergent regulatory features of trichome development in poplar, which produces long, unbranched trichomes (Plett et al. 2010; Feodorova and Alexandrov 2020). Although non-glandular trichomes have been comparatively understudied relative to the glandular trichomes as specialized chemical factories (Huchelmann, Boutry, and Hachez 2017), recent work has demonstrated that poplar trichomes are also capable of terpenoid production (Bewg *et al*., 2022).

The poplar 717 single cell atlas will enable data mining by the community at single cell resolution. Through exploration of the atlas, researchers can investigate cell type expression profile of their genes of interest, shortlist spatiotemporally restricted genes and their regulatory regions, investigate the cell-type specificity of biological processes of interest, explore the diversity of cell types shared across different organs, and integrate this atlas with data from other experiments to explore new hypotheses. We believe this atlas represents an important step forward in tree research by providing a comprehensive single-cell atlas and establishing a valuable resource for tree synthetic biology. As shown in our deep analysis of the epidermis, the atlas enabled discovery of cell type specific gene expression for trichomes that can be exploited in metabolic engineering in the future.

## Materials and Methods

### Plant Growth Conditions and Sampling

Plant material was sampled from female interspecific hybrid aspen *Populus tremula* x *P. alba* INRA 717-1B4 (Leple et al. 1992). All plants sampled for the scRNA-seq atlas were grown in Conviron PG40 growth chambers under 24 °C day/18 °C night cycle, with a day length of 18 hours, 500 µmol/m^2^/s light with100% far red supplement. The height ranged from 92 to 150 cm at sampling. For leaves, three developmental stages were sampled starting with the first fully open leaf larger than 2 cm as leaf 1 (L1) and sampling downwards for leaf 3 (L3) and leaf 5 (L5). For stems, the first internode to the third internode (as denoted by leaf number) were collected as primary stem (PS), and from internode five to seven was collected as secondary stem (SS). Shoot apical meristems (SAMs) 1 cm in length were collected from plants with only one stem; leaf primordia were removed. Samples were collected between 9:00-10:00 am, 3-4 hours after dawn. Samples were pooled with 13 L1, 10 L3, and 4 L5 leaves per pool per each of two replicates. For stem, 14 primary and 4 secondary stem sections were pooled per each of two replicates. For SAM, 12 apices were pooled per each of two replicates.

The glabrous poplar 717 mutants KO-2, KO-25 and KO-46 (Bewg et al. 2022), along with wild-type plants were propagated by single-node cuttings and grown in a greenhouse as previously described (Frost et al. 2012). Newly emerged L1 leaves were sampled for bulk RNA-seq between 11:30am - 12:00pm. One replicate each of three independent mutant events or three replicates for poplar 717 were collected.

### Protoplast Isolation and Single Cell Library Preparation

Each organ and developmental stage was processed independently using an optimized protoplast isolation protocol. Before immersing samples in an enzymatic solution, leaves were cut into ∼1 mm slices, and apical meristems (1 cm in length) were trimmed of leaf primordia and bracts and sectioned into 4 longitudinal portions (in half and then each half in half again). For both PS and SS, outer stem (bark) was separated via peeling from the inner stem (Lin et al. 2014; Tung et al. 2023) prior to immersion in the enzymatic solution. Protoplasts were isolated as previously described (C. Li et al. 2023). For all samples except SAM, which required adjusted conditions, samples were immersed in the enzymatic solution and vacuum infiltrated for 10 minutes at 400 mbar. Samples were then placed into plastic petri dishes and shaken at room temperature for 1 hour 45 minutes at 50 rpm and for 5 minutes at 80-100 rpm to facilitate protoplast detachment. Protoplasts were filtered through 70 μm and 40 μm mesh, then 5 mL of storage solution (0.4 M mannitol, 20 mM MES, 0.20% 2-mercaptoethanol, 1 mM calcium chloride, 0.10% BSA, pH 5.7) were used to rinse the sample on the petri dish to increase protoplast recovery. Protoplasts were then pelleted by spinning the filtrate at 100 x *g* for 5 minutes at 4 °C. The supernatant solution was discarded, and protoplasts were gently resuspended in 2 mL storage solution. An OptiPrep (SigmaAldrich, St. Louis, MO. Cat. No. D1556) (Graham 2002) density gradient cleanup was performed to separate protoplasts from any leftover debris.

ScRNA-seq libraries were constructed using Fluent Biosciences PIP-seq T20 3’ Single Cell RNA kits (v4.0Plus) (Clark et al. 2023) according to manufacturer’s instructions. A total of ∼16,200 (L1), ∼23,400 (L3), ∼13,500 (L5), ∼40,000 (SS), and ∼39,600 (SAM) cells were used as input for library preparation for each of two replicates. For primary stem (PS), ∼25,200 (first replicate) and ∼22,500 (second replicate) cells were used. Sequencing was performed by the DNA Technologies and Expression Analysis Cores at the UC Davis Genome Center (RRID: SCR_017740) on an Element Biosciences AVITI (San Diego, CA).

### Single Cell Data Processing

PIPseeker barcode (v3.3) was used to clean read 2 according to the instruction manual (PIPseeker v3.1 User Guide). Reads were then aligned onto the haplotype resolved 717 genome (R. Zhou et al. 2023) and its organellar genomes (chloroplast and mitochondrion) (Kersten et al. 2016) using STARsolo (Kaminow, Yunusov, and Dobin 2021) with the following parameters: soloType CB_UMI_Simple, --soloCBwhitelist (using the barcode whitelist generated by PIPseeker), --soloUMIlen 12, --soloBarcodeReadLength 0, --alignIntronMax 5000, --soloFeatures GeneFull, --soloMultiMappers EM. After mapping, the EmptyDrops_CR cell filtering algorithm was run on the output to count multimapping reads, addressing high multimapping rates due to the haplotype resolved reference genome.

### Single-Cell RNA-Seq Data Analysis

Expression matrices were loaded into R v4.3.2 and analyzed using Seurat v5.2.1 (Stuart et al. 2019) using standard pre-processing workflows. Cells containing more than 5% organellar reads were excluded from the analysis, as well as cells containing less than 500 or more than 10,000 features, and/or less than 500 or more than 30,000 UMIs. Cell filtering cutoff thresholds based on the number of RNA features were determined on a case-by-case basis. The data was then log normalized and the 3,000 most variable genes were selected for principal components analysis (PCA), which informed the number of PCs to be used for dimensionality reduction via UMAP (Uniform Manifold Approximation and Projection). DoubletFinder (McGinnis, Murrow, and Gartner 2019) was used to detect and exclude doublets from the dataset.

Using the Seurat *SelectIntegrationFeatures()* function, 3,000 features were selected. Integration anchors were found using the Seurat *FindIntegrationAnchors()* function and the “rpca” (reciprocal PCA) reduction method. The two replicates for each combination of organ and developmental stage were integrated using the Seurat *IntegrateData()* function. All datasets were then merged to generate a single Seurat object using the *merge()* function. To annotate the data, a set of marker genes was curated.

### Leaf Bulk RNA Isolation and Sequencing

Leaves were ground with a mortar and pestle in liquid nitrogen and RNA was isolated using a modified hot borate method (Wan and Wilkins 1994; Wood et al. 2026). RNA was treated with DNase using the Turbo DNase Free Kit (ThermoFisher Scientific, Waltham, MA). RNA-seq libraries were constructed at the Texas A&M AgriLife Research Genomics and Bioinformatics Service using the PerkinElmer NEXTFLEX Rapid Directional RNA-Seq Kit 2.0. Subsequent libraries were sequenced on a NovaSeq X Plus to 150 nt in paired end mode (Illumina, San Diego, CA).

### Bulk RNA-Seq Data Processing

Adapters and low-quality sequence were cleaned from the reads using Cutadapt v4.9 (Martin 2011) using the parameters; -q 30 -m 75 --trim-n -n 2. Cleaned reads were then used to calculate Transcripts per Million (TPM) values for each gene using Kallisto v0.48.0 (Bray et al. 2016) in stranded mode using the primary transcripts from the 717 v5 haplotype-resolved assembly (R. Zhou et al. 2023). Cleaned reads were also aligned to the 717 v5 haplotype resolved assembly using HISAT2 v2.2.1 (Kim et al. 2019) in stranded mode with a max intron size of 5kb and output formatted for Cufflinks. The alignments were then counted using HTSeq v2.0.2 (Anders, Pyl, and Huber 2015) with the parameters: --minaqual=1 --type=gene --mode=union --nonunique all. The parameters --nonunique all and a --minaqual=1 were set to allow for multimapping reads (MAPQ = 1) from HISAT2 to be counted.

### Bulk RNA-Seq Differential Gene Expression Analysis

The DESeq2 1.44.0 (Love, Huber, and Anders 2014) package in R was used to perform differential gene expression, of the mutant pool (KO-2, KO-25 and KO-46) vs wild-type poplar 717. The “contrasts” argument was used to test if the log_2_ fold change was equal to 0. Results were filtered to only contain differentially expressed genes that had a log_2_ fold-change threshold of ±2, and an adjusted p-value ≤ 0.05. To restrain high log_2_ fold changes associated with genes that had low expression values we used the *lfcShrink()* function within DESeq2.

### Phylogenetic Analysis

Poplar 717 orthologs of *Arabidopsis* CSL genes were identified via Orthofinder v2.5.5 (Emms and Kelly 2019). The CESA and CSL phylogeny was generated by aligning representative gene models with MAFFT (v7.526) (Katoh and Standley 2013) using the option “--localpair”. The phylogeny was built with RAXML-NG (v.1.2.2) (Kozlov et al. 2019) using model LG+G and 1,000 bootstrap replicates and visualized using TVBOT (Xie et al. 2023).

### MATL16 promoter cloning, transgenic characterization, and microscopy

An 1,844 bp *PtMATL16* (PtXaTreH.10G194200) promoter fragment was amplified from poplar 717 genomic DNA isolated from young leaves using a plant DNA minipreparation protocol (Dellaporta, Wood, and Hicks 1983). The first PCR was performed using the forward primer MATP.F1 (5′-attcgagctctcccatatg<u>GTCGAC</u>TTGACGGGATCAATCTTTATAGT-3′) and the reverse primer MATP.R1 (5′-GGGAGAGTGAACTCAGTGTTT-3′). A second, nested PCR was carried out using the same forward primer and the nested reverse primer MAT.R2. (5′-cttcgccactgctcaccat<u>ACTAGT</u>GGATGAGAAGCTAGAAGTCGTAA-3′), with 10-fold diluted primary PCR products as the template. The nested PCR products, containing vector homology arms (lowercase in primer sequences) and restriction sites (underlined), were cloned into *pH35S::fuGFP* at the *Sal*I and *Spe*I sites to replace the CaMV 35S promoter using the NEBuilder HiFi DNA Assembly Cloning Kit (New England Biolabs). The *pH35S::fuGFP* vector (kindly provided by Maria Ortega) was derived from *pH35S::mEGFP* (Behrendorff, Borràs-Gas, and Pribil 2020) (Addgene plasmid #135321) by replacing *mEGFP* with *fuGFP* (Moratti et al. 2024) at the *Spe*I and *Eco*RI sites. The resulting *PtMATL16::fuGFP* construct was confirmed by Sanger sequencing and transformed into *Agrobacterium tumefaciens* strain C58pMP90 using the freeze-thaw method (Höfgen and Willmitzer 1988).

Poplar 717 transformation was performed as previously described (Ortega et al. 2023). Regenerated plants were transplanted to soil and maintained in a walk-in growth room. Three independent transgenic events and multiple WT plants were examined by confocal microscopy. Leaves from two-month-old plants were cut into 5-8 mm² segments, mounted on microscope slides, and gently covered with a coverslip using a paintbrush to avoid trichome damage. Fluorescence imaging was performed using an LSM 880 laser scanning confocal microscope (Carl Zeiss, Oberkochen, Germany) with excitation at 488 nm (argon ion laser) and emission collected between 493 and 598 nm. Images were acquired as single optical sections or as z-stack series and processed using FIJI (Image J) (Schindelin et al. 2012).

## Data Availability

Raw sequencing data are available at the National Center for Biotechnology Information Sequence Read Archive BioProject PRJNA1401748 (to be made public upon acceptance). We created an interactive web viewer for the poplar shoot atlas using the Streamlit Python library (http://bio-poplar-atlas.com). The processed scRNA-seq datasets were converted to h5ad format, and Scanpy was used for visualization on the viewer.

## Acknowledgements

The authors acknowledge Gilles Pilate (Institut National de la Recherche Agronomique, France) for providing poplar clone INRA 717-1B4, Maria Ortega for providing pH35S::mEGFP, and Muthugapatti Kandasamy at the Biomedical Microscopy Core, University of Georgia, for confocal imaging assistance. This material is based upon work supported by the U.S. Department of Energy, Office of Science, Office of Biological and Environmental Research program under Award Number DE-SC0023338.

